# Reproducible Computational Workflows with Continuous Analysis

**DOI:** 10.1101/056473

**Authors:** Brett K. Beaulieu-Jones, Casey S. Greene

**Affiliations:** Genomics and Computational Biology Graduate Group, Perelman School of Medicine, University of Pennsylvania.; Department of Systems Pharmacology and Translational Therapeutics, Perelman School of Medicine, University of Pennsylvania

## Abstract

Reproducing experiments is vital to science. Being able to replicate, validate and extend previous work also speeds new research projects. Reproducing computational biology experiments, which are scripted, should be straightforward. But reproducing such work remains challenging and time consuming. In the ideal world we would be able to quickly and easily rewind to the precise computing environment where results were generated. We would then be able to reproduce the original analysis or perform new analyses. We introduce a process termed “continuous analysis” which provides inherent reproducibility to computational research at a minimal cost to the researcher. Continuous analysis combines Docker, a container service similar to virtual machines, with continuous integration, a popular software development technique, to automatically re-run computational analysis whenever relevant changes are made to the source code. This allows results to be reproduced quickly, accurately and without needing to contact the original authors. Continuous analysis also provides an audit trail for analyses that use data with sharing restrictions. This allows reviewers, editors, and readers to verify reproducibility without manually downloading and rerunning any code. Example configurations are available at our online repository (https://github.com/greenelab/continuous_analysis).

## The Current State of Reproducibility

Leading scientific journals have targeted reproducibility to increase readers’ confidence in results and reduce retractions^1–5^. In a recent survey, 90% of researchers acknowledged a reproducibility crisis^6^. Research that uses computational protocols should be particularly amenable to reproducible workflows because all of the steps are scripted into a machine-readable format. But written descriptions of computational approaches can be difficult to understand and may lack required parameters. Even when results can be reproduced, the process often requires a substantial time investment and help from the original authors. Garijo et al.^7^ estimated it would take 280 hours for a non-expert to reproduce a paper describing a computational construction of a drug-target network for *Mycobacterium tuberculosis*^8^. These are the good scenarios: the results behind most computational papers are not readily reproducible^7,9–11^.

The practice of “open science” has been discussed as a means to aid reproducibility^3,12^. In open science the data and source code are shared. Sharing can also extend to intermediate results and project planning^13^. Sharing data and source code is currently necessary but not sufficient to make research reproducible. Even when code and data are shared, it remains difficult to reproduce results due to differing computing environments, operating systems, library dependencies etc. It is common to use one or more open source libraries on a project, and research code quickly becomes dependent on old versions of these libraries as software advances^14^. These old or broken dependencies make it difficult for readers and reviewers to recreate the environment of the original researchers, whether to validate or extend their work.

An example of where sharing data does not automatically make science reproducible occurs in the most standard of places: differential gene expression analysis. Such analyses are routine. Our understanding of the genome, including transcriptome annotations, have improved and updated probe set definitions are available^15^. Analyses relying on unspecified probe set definitions cannot be reproduced using current definitions.

We analyzed the fifteen most recently published papers that cite Dai et al., a common source for custom chip description files (CustomCDF), that were accessible at our institution^16–31^. We identified these manuscripts using Web of Science on May 31, 2016. We recorded the number of papers that cited a version of CustomCDF, as well as which version was cited. We expect this analysis to provide an upper bound on reproducible work: these papers specifically cited the source of their CDFs. Of these fifteen papers, nine (60%) specified which version they used. These nine used versions 11, 15, 16, 17, 18, and 19 of the BrainArray CustomCDF.

This initial analysis was performed based on article recency without regard to article impact. To determine the extent to which this issue affects high impact papers, we performed a parallel evaluation for the ten most cited papers^32–41^ that cite Dai et al. We determined the ten most cited papers using Web of Science on May 31, 2016. Of these ten papers, one^38^ (10%) specified which version of the CustomCDF was used. That paper used version 11 of the BrainArray CustomCDF.

We sought to determine which versions were currently in use in the field. We asked three individuals who performed microarray analysis recently, and we accessed and evaluated two cluster systems used for processing data. We found that each individual had one of the three most recently released versions installed (18, 19, and 20), and versions 18 and 19 were installed on cluster systems.

To evaluate the impact of differing CDF versions, we downloaded a recently published public gene expression dataset (GEO Series Ascension number GSE47664). This experiment examined differential expression between normal HeLa cells and HeLa cells with TIA1 and TIAR knocked down^42^. We performed a parallel analysis using each of the three versions that we found installed on machines that we could access (18, 19, and 20). Each version identifies a different number of significantly altered genes (Figure 1A), demonstrating the challenge of reproducible analysis. We simulated a parallel analysis of differential expression using Docker containers on mismatched machines^43^. This specifies the CDF version and produces the same number and set of differentially expressed genes for a given version across machines (v18 example in Figure 1B). Had continuous analysis been used for papers citing the BrainArray CustomCDF their computational results would be easily replicated.

**Figure 1.**
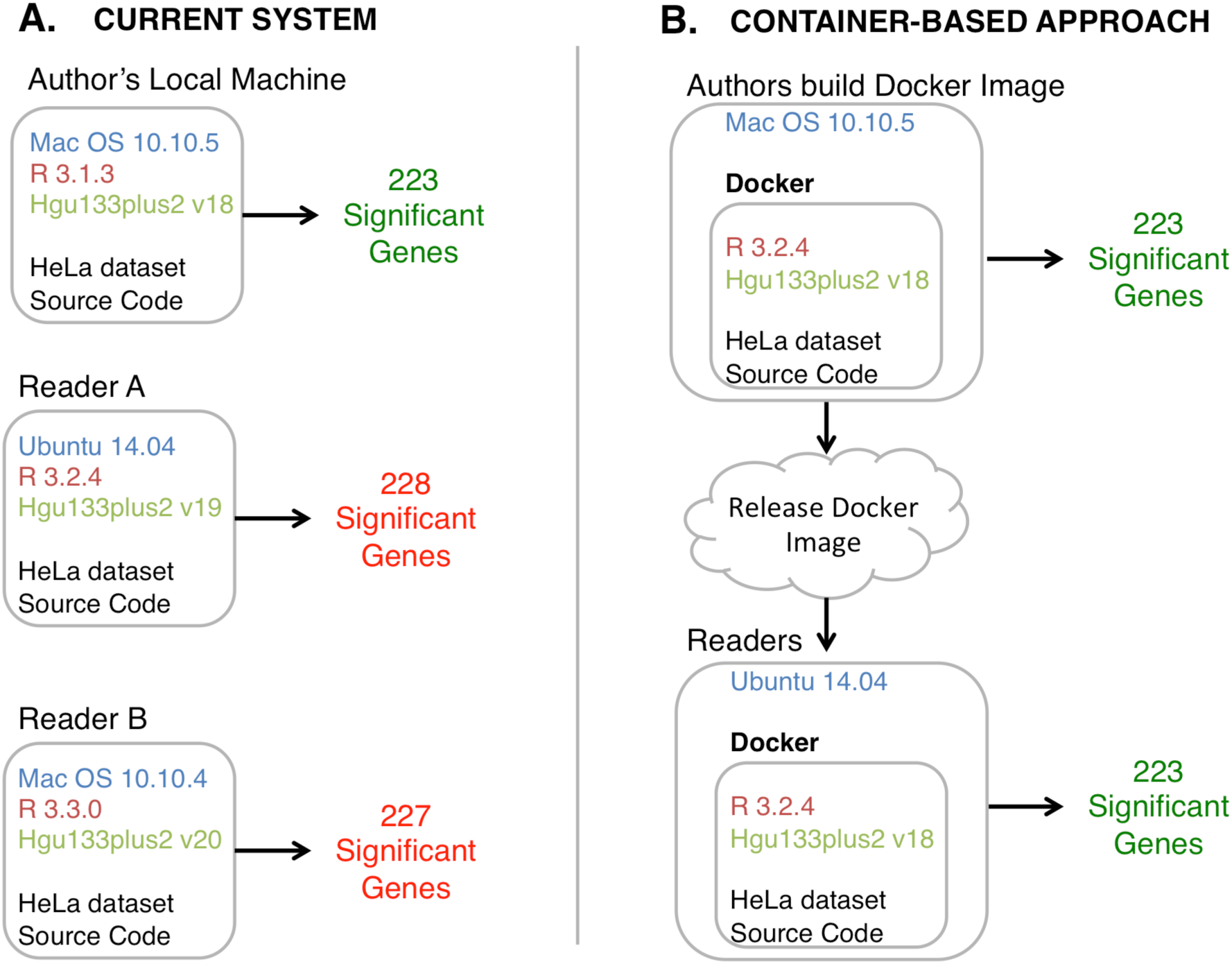
Current state of research computing vs. container-based approaches. **A.)** The status quo requires a reader or reviewer to find and install specific versions of dependencies. These dependencies can become difficult to find and may become incompatible with newer versions of other software packages. Different versions of packages identify different numbers of significantly differentially expressed genes from the same source code and data. **B.)** Containers define a computing environment that captures dependencies. In container-based systems, the results are the same regardless of the host system.

## Continuous Analysis in Computational Workflows

We developed continuous analysis to produce a verifiable end-to-end run of computational research with minimal start-up costs. In contrast with the status quo, continuous analysis preserves the computing environment and maintains the versions of dependencies. We described the benefits of containerized approaches above, but maintaining, running and distributing Docker images manually would become time consuming. Integrating Docker into a continuous scientific analysis pipeline meets three criteria: (1) anyone can re-run code in a computing environment matching the original authors (Supplemental Figure 1); (2) readers and reviewers can follow exactly what was done in an “audit” fashion without running code (Supplemental Figure 2 & 3); and (3) the solution imposes zero to minimal cost in terms of time and money on the researcher, depending on their current research process.

Continuous analysis extends continuous integration^44^, a common practice in software development and deployment. Continuous integration is a software development workflow that triggers an automated build process whenever developers check their code in to a source control repository. This automated build process runs test scripts if they exist. These tests can catch bugs introduced into software. Software that passes tests is automatically deployed to remote servers.

For continuous analysis (Figure 2), we repurpose these services in order to run computational analyses, update figures, and publish changes to online repositories whenever relevant changes are made to the source code. When an author is ready to release code or publish their work they can export the most recent continuous integration run. Because this process generates results in a clean and clearly defined computing environment without manual intervention, reviewers can be confident that the analyses are reproducible.

**Figure 2.**
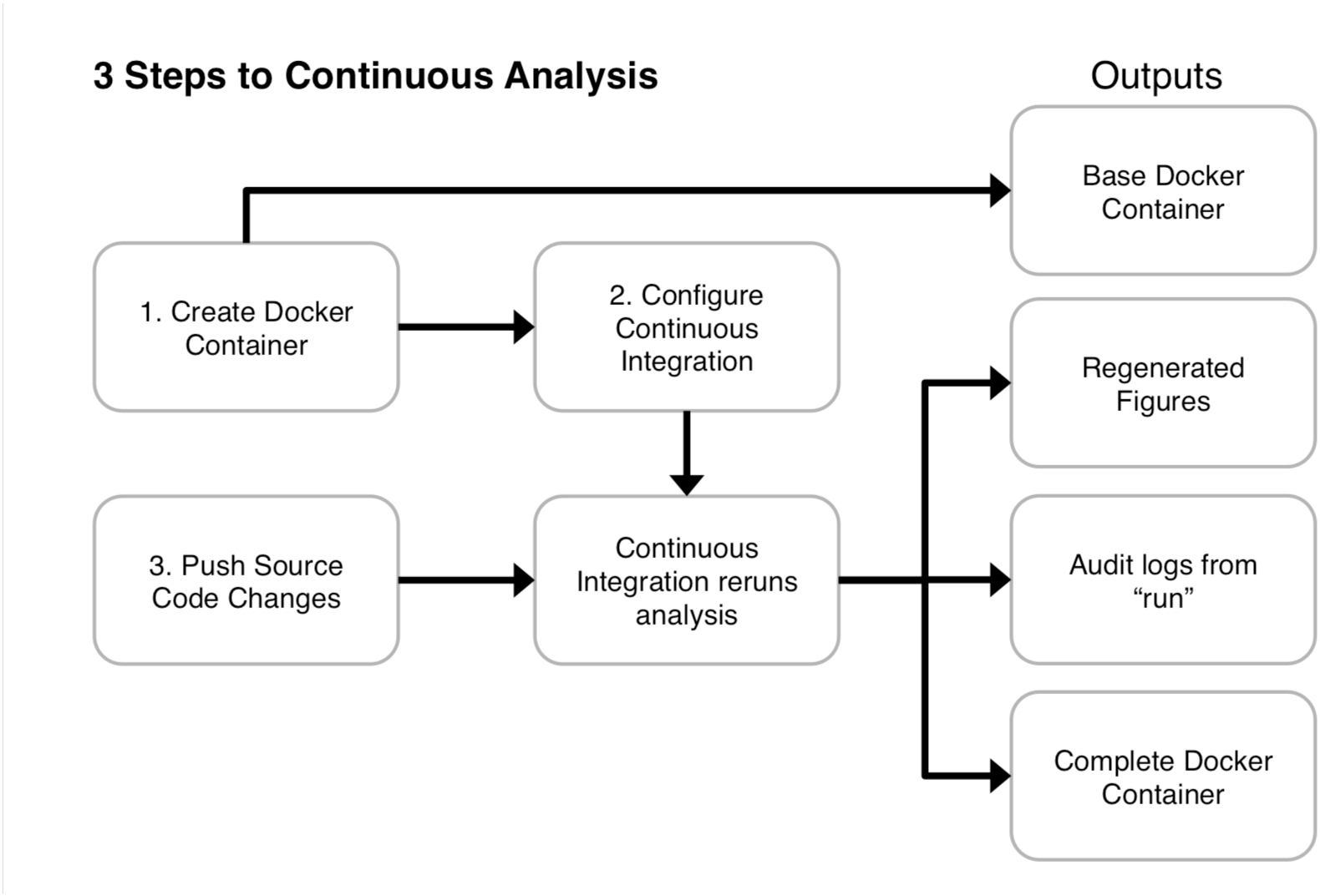
Continuous analysis can be set up in three primary steps (numbered 1, 2, and 3). (1) The researcher creates a Docker container with the required software. (2) The researcher configures a continuous integration service to use this Docker image. (3) The researcher pushes code that includes a script capable of running the analyses from start to finish. The continuous integration provider runs the latest version of code in the specified Docker environment without manual intervention. This generates a Docker container with intermediate results that allows anyone to rerun analysis in the same environment, produces updated figures, and stores logs describing everything that occurred. Example configurations are available in the supplementary materials as well as our online repository (https://github.com/greenelab/continuous_analysis). Because code is run in an independent, reproducible computing environment and produces detailed logs of what was executed, this practice reduces or eliminates the need for reviewers to re-run code to verify reproducibility.

In each project we maintain dependencies with the free open-source software tool Docker^45^. Docker defines an “image” that allows users to download and run a container, a minimalist virtual machine with a predefined computing environment. Docker images can be several gigabytes in size, but once downloaded can be started in a matter of seconds and has minimal overhead^14^. In addition, Docker images can be easily tagged to coincide with software releases and paper revisions. At the time of submission, authors can run the `docker savè command to export a static file that can be uploaded to services such as Figshare or Zenodo to receive a DOI. For example, we have uploaded our continuous analysis environment for the examples in this paper^46^.

To set up continuous analysis, a researcher needs to do three things. First they must create a Dockerfile, which specifies a list of dependencies. Second, they need to connect a continuous integration service to their version control system and provide the commands to run their analysis. Finally, they need to commit and push changes to their version control system. Many researchers already perform the first and third tasks in their standard workflow.

The continuous integration system will automatically rerun the specified analysis with each change, precisely matching the source code and results. It can also be set to listen and run only when changes are marked in a specific way, e.g. by committing to a specific ‘staging’ branch. For the first project, this process can be put into place in less than a day. For subsequent projects, this can be done in under an hour.

## Setting up Continuous Analysis

We have created a GitHub repository with instructions for paid, local, and cloud-based continuous analysis setups^47^. These are fully detailed in the supplementary materials and online repository. Here we describe how continuous analysis can be setup using the free and open source Drone software on a researcher’s personal computer and connected to the GitHub version control service. This setup is free to users.

1. Install Docker on the computer.
2. Pull the Drone image via docker:

> sudo docker pull drone/drone:latest
3. Create a new application in GitHub (Figure 3).
4. Add a webhook to the GitHub project (Figure 4). This will notify the continuous integration server of any updates pushed to the repository.
5. Create a configuration file on the Drone computer at /etc/drone/dronerc filling in the client information provided by GitHub

> REMOTE_DRIVER=github
>
> REMOTE_CONFIG=https://github.com?client_id=….&client_secret=….
6. Run the drone container

> sudo docker run drone/drone:latest

**Figure 3.**
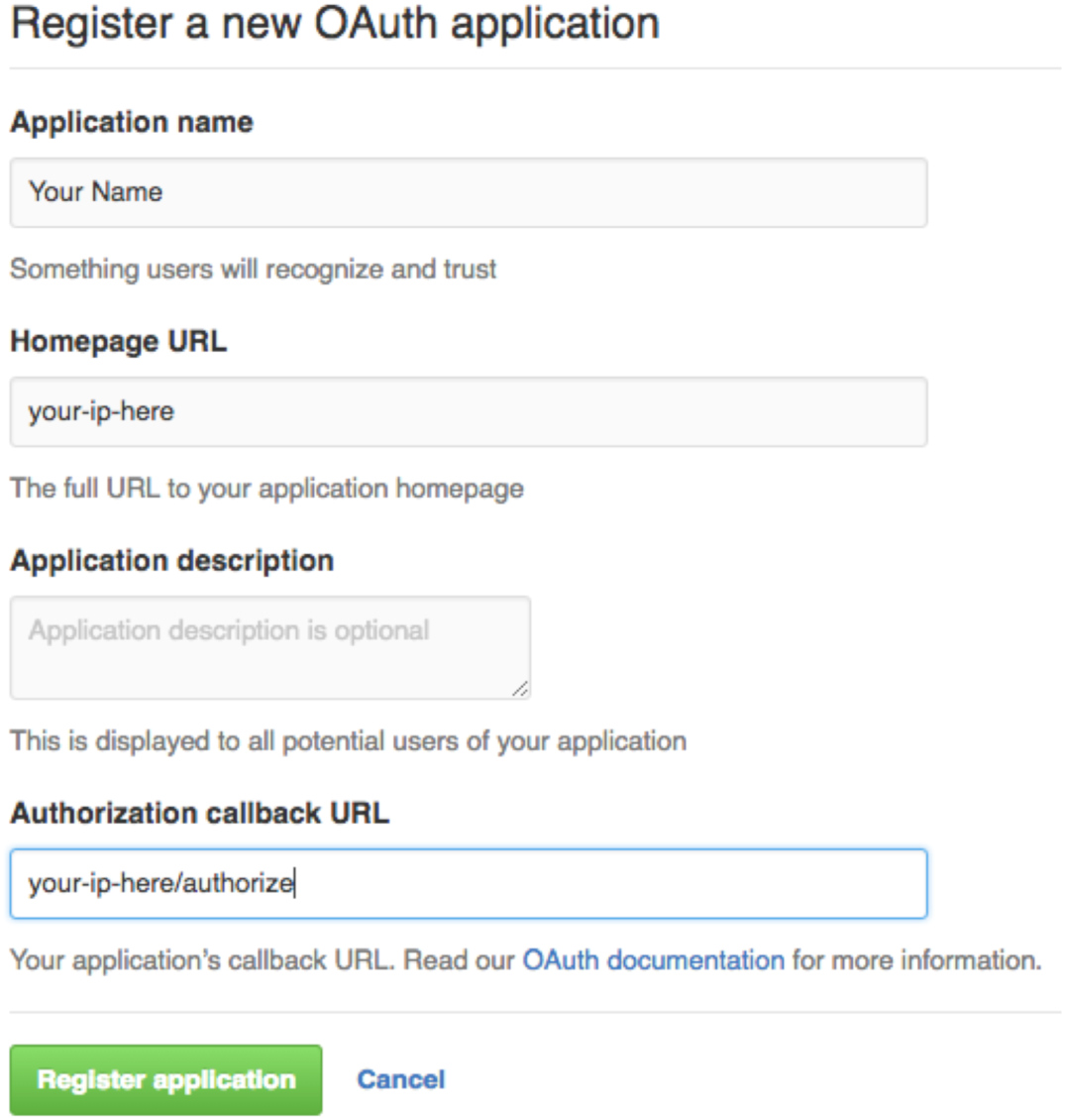
Register a new application for the Drone continuous integration server. Set the homepage URL to be the IP address of the Drone computer. Set the callback URL to the same IP address followed by /authorize.

**Figure 4.**
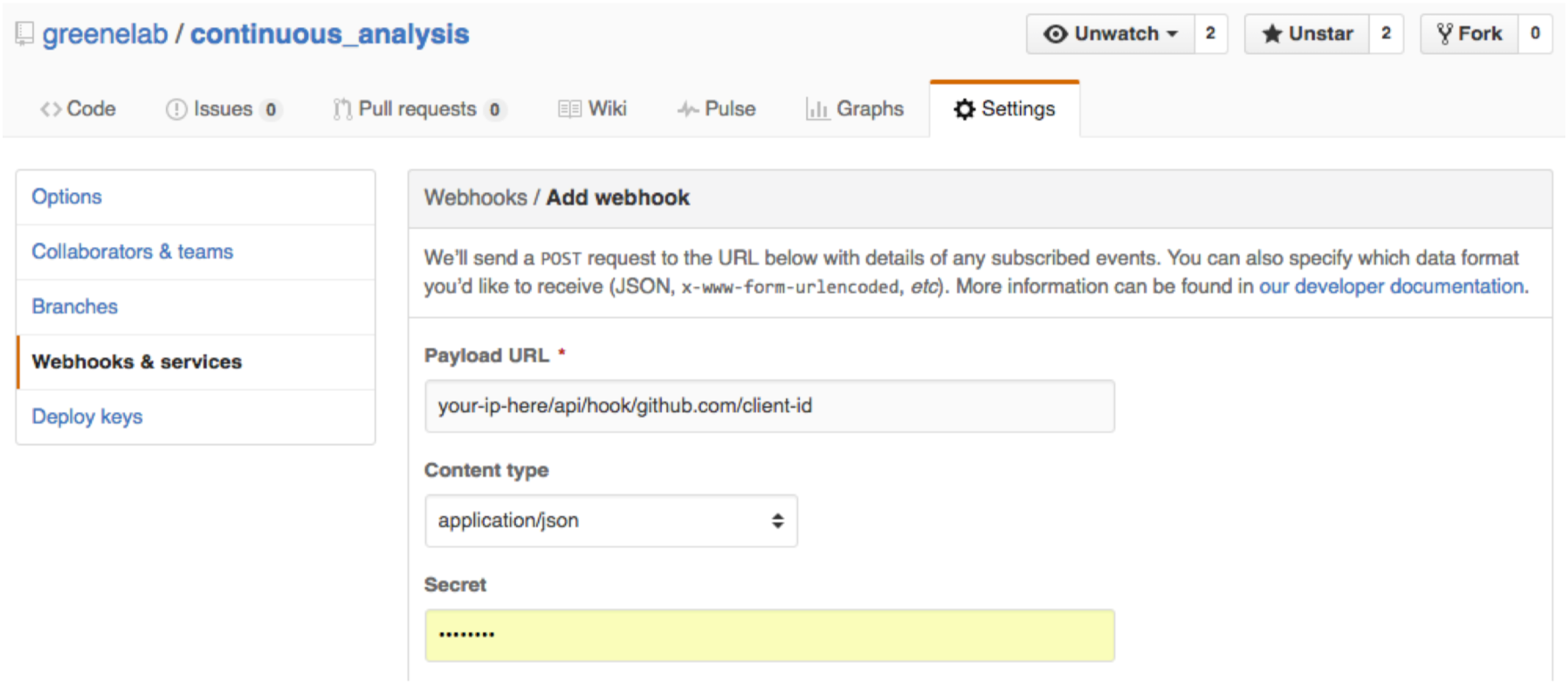
Register a new application for the Drone continuous integration server. The payload URL should be in the format of your-ip/api/hook/github.com/client-id

Continuous analysis can be performed with dozens of full service providers or a private installation on a local machine, cluster or cloud service^47^. Full service providers can be set up in minutes but may have computational resource limits or monthly fees. Private installations require configuration but can scale to a local cluster or cloud service to match the computational complexity of all walks of research. With free, open-source continuous integration software^48^, computing resources are the only associated costs.

## Using Continuous Analysis

After setup, running continuous analysis is simple and fits into existing research workflows that use source control systems. We have used continuous analysis in our own work^49^. We have also prepared three example repositories (detailed in supplemental materials):

1. An example demonstrating the setup of continuous analysis with a wide variety of services and configurations (highlighted below).
2. An easy to follow basic phylogeny tree building example, combining sequence alignment using MAFFT^50^, format conversion using EMBOSS Seqret^51^, and tree calculation and drawing using.
3. An RNA expression analysis workflow examining organoid models of pancreatic cancer in mice based on work from Boj et al.^52^ using details and source code published by Balli^53^. This example shows the ability of continuous analysis to scale to large computations. This example uses kallisto^54^, limma^55,56^, and sleuth^57^ to analyze 150GB of gene expression data and approximately 480 million reads.

To demonstrate the setup process and different configurations of continuous analysis we show a simple example of continuous analysis with kallisto. The recently published software tool kallisto quantifies transcript abundance in RNA-seq data. Our example re-runs the examples provided in kallisto with each commit to a repository.

1. Add a script file to re-run custom analysis. For Drone, this is a .drone.yml file that specifies commands to run each step of the analysis. An example configuration is available in the continuous analysis GitHub repository as well as the supplemental materials.
2. Commit changes to the source control repository.
3. Push changes to GitHub.

The configured continuous integration service automatically runs the specified script. We configured this to rerun the analysis, regenerate the figures, and commit updated versions to the repository. The service provides a complete audit log of what was run in the clean continuous integration environment (Figure 5). By generating and pushing updated figures, this process also generates a complete change log for each result (Figure 6). Interactive development tools, such as Jupyter^58,59^, RMarkdown^60^,61 and Sweave^62^ can be incorporated to present the code and analysis in a logical graphical manner. For example, we recently used Jupyter with continuous analysis in our own publication^63^ and corresponding repository^49^.

**Figure 5.**
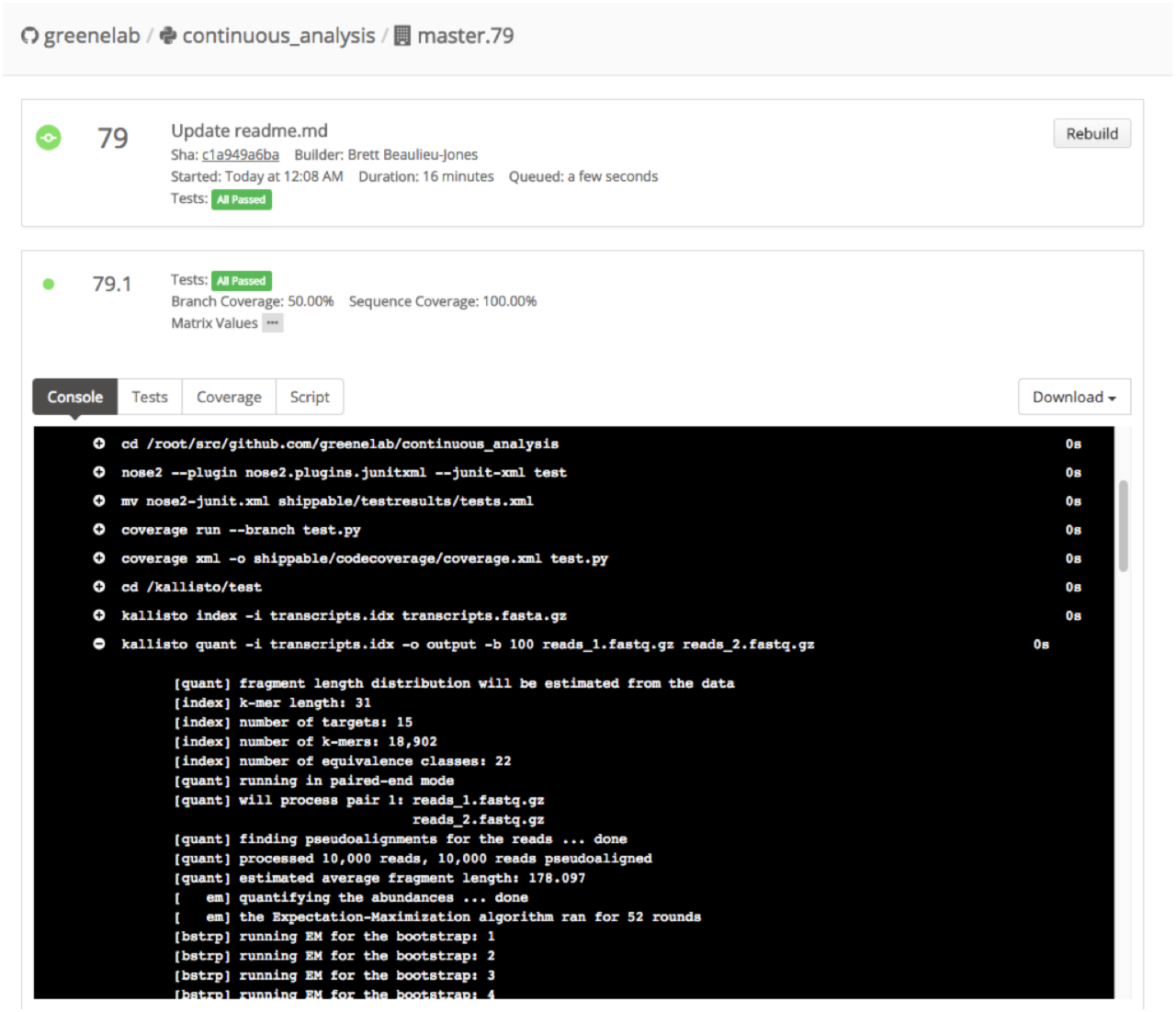
Audit logs from a continuous integration run with the service Shippable for the kallisto example.

**Figure 6.**
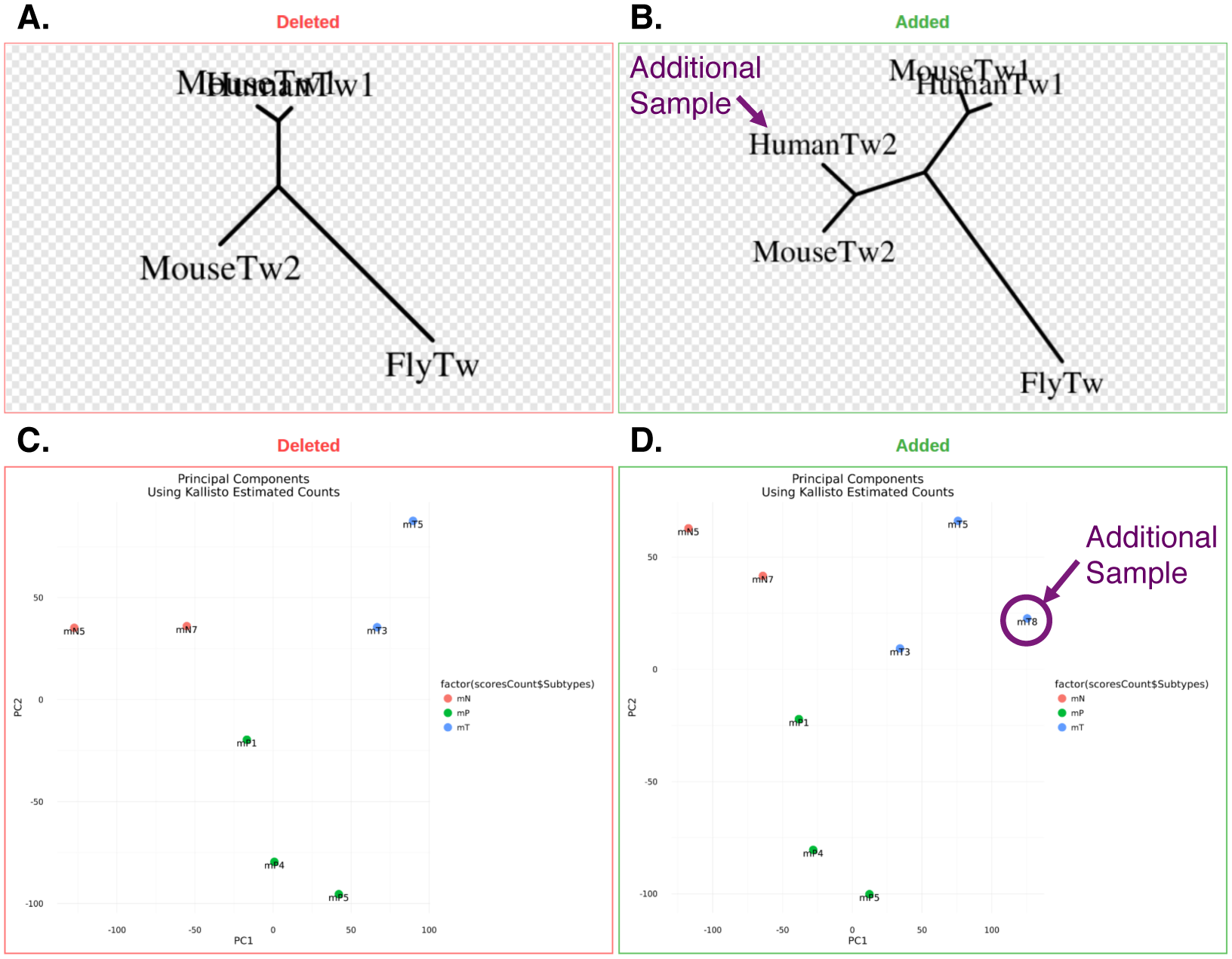
Resulting figures from the run are committed back to Github where changes between runs can be viewed. **A.)** The effect of adding an additional gene (HumanTw2) to a phylogenetic tree-building example. **B.)** The effect of adding an additional gene (mt8) to an RNA-seq differential expression experiment PCA plot.

In summary, continuous analysis provides the results of a verifiable end-to-end run in a “clean” environment. Because continuous analysis runs automatically in the background, no transition is needed between the exploration and publication phases of a scientific project. The audit trail provided by continuous analysis allows reviewers and editors to provide sound judgment on reproducibility without a large time commitment. If readers or reviewers would like to re-run the code on their own (e.g. to change a parameter and evaluate the impact on results), they can easily do so with the Docker container containing the final computing environment and intermediate results. Version control systems provide the capability to watch for updates. Readers can “star” or “watch” a repository on services such as Github, Gitlab, and Bitbucket to be automatically notified of changes and updated runs. Wide adoption of these systems throughout the publication process could allow reviewers and editors to automatically be notified of updated results.

## Continuous analysis provides an audit trail for reproducible analyses of closed data

Continuous analysis can be even more powerful when working with closed data that cannot be released. Without continuous analysis, reproducing computational analyses based on closed data is dependent on the original authors completely and exactly describing each step, a process that may be an afterthought and relegated to extended methods. Readers must then diligently follow complex written instructions without intermediate confirmation they are on the right track. The containers produced during continuous analysis include a matching environment for replication as well as intermediate results. This allows readers to determine where their results diverge from the original work and to determine whether divergence is due to software-based or data-based differences.

## Best practices with continuous analysis

We suggest a development workflow where continuous analysis runs only on a single branch (Supplemental Figure 4). Researchers can push to this branch when they believe they are ready for a release to avoid running the full process during incomplete updates. If the updates to this branch succeed, the changes are then automatically carried over to the master or production branch and released. We recommend exporting both the before and after processing Docker images and uploading to an archival service like Figshare or Zenodo. The archived images can then be cited to guide readers to the version used in the manuscript^46^. For convenience, the images can also be shared through the Docker Hub registry.

It may currently be impractical to use continuous analysis for generic preprocessing steps involving very large data or analyses requiring particularly high computational costs. In particular, steps that take days to run or incur substantial costs in computational resources may not be amenable with existing providers^64^. One day, continuous analysis systems specifically designed for scientific workflows may facilitate reproducible workflows in these settings. For now, researchers may need to use discretion when preprocessing via continuous analysis, as it may be computationally intractable to reanalyze after each commit to a staging branch. Researchers may elect to run only the final workflow through this process, or may elect to employ continuous analysis after standard but computationally expensive preprocessing steps are completed.

For small datasets and less intensive computational workflows it is easiest to use a full service continuous integration service. These services have the smallest setup times. With private data or when data size and computational complexity scale it becomes necessary to setup a local privately hosted continuous integration server. Cluster or cloud based continuous integration servers can handle the largest workflows.

## The impact of reproducible computational research

Reproducibility can have wide-reaching benefits for the advancement of science. For authors, easily reproducible work is a sign of quality and credibility. Continuous analysis addresses the reproducibility of computationally analyses in the narrow sense: generating the same results from the same inputs. It does not solve reproducibility in the broader sense: how robust results are to parameter settings, starting conditions and partitions in the data. Continuous analysis lays the groundwork needed to address reproducibility and robustness of findings in the broad sense.

## Acknowledgements

This work was supported by the Gordon and Betty Moore Foundation under a Data Driven Discovery Investigator Award to CSG (GBMF 4552) and supported by a Commonwealth Universal Research Enhancement (CURE) Program grant from the Pennsylvania Department of Health. We would like to thank David Balli for providing the RNA-seq analysis design, Katie Siewert for providing the phylogenetic analysis design, and Alex Whan for contributing a Travis-CI implementation.

## References

1. Rebooting review. Nat Biotech. 2015;33(4): 319. http://dx.doi.org/10.1038/nbt.3202.

2. Software with impact. Nat Meth. 2014;11(3): 211. http://dx.doi.org/10.1038/nmeth.2880.

3. Peng RD. Reproducible Research in Computational Science. Science (80-). 2011;334(6060): 1226–1227. doi:10.1126/science.1213847.

4. McNutt M. Reproducibility. Science (80-). 2014;343(6168): 229. http://science.sciencemag.org/content/343/6168/229.abstract.

5. Illuminating the black box. Nature. 2006;442(7098): 1. http://dx.doi.org/10.1038/442001a.

6. Baker M. 1,500 scientists lift the lid on reproducibility. Nature. 2016;533(7604): 452–454. doi:10.1038/533452a.

7. Garijo D, Kinnings S, Xie LL, et al. Quantifying reproducibility in computational biology: The case of the tuberculosis drugome. PLoS One. 2013;8(11). doi:10.1371/journal.pone.0080278.

8. Kinnings SL, Xie LL, Fung KH, Jackson RM, Xie LL, Bourne PE. The Mycobacterium tuberculosis drugome and its polypharmacological implications. PLoS Comput Biol. 2010;6(11). doi:10.1371/journal.pcbi.1000976.

9. Bell AW, Deutsch EW, Au CE, et al. A HUPO test sample study reveals common problems in mass spectrometry-based proteomics. Nat Methods. 2009;6(6): 423–430. doi:10.1038/nmeth.1333.

10. Ioannidis JPA, Allison DB, Ball CA, et al. Repeatability of published microarray gene expression analyses. Nat Genet. 2009;41(2): 149–155. doi:10.1038/ng.295.

11. Hothorn T, Leisch F. Case studies in reproducibility. Brief Bioinform. 2011;12(3): 288–300. doi:10.1093/bib/bbq084.

12. Groves T, Godlee F. Open science and reproducible research. BMJ. 2012;344. doi:10.1136/bmj.e4383.

13. ThinkLab. https://thinklab.com/. Accessed January 1, 2016.

14. Boettiger C. An introduction to Docker for reproducible research, with examples from the R environment. ACM SIGOPS Oper Syst Rev Spec Issue Repeatability Shar Exp Artifacts. 2015;49(1): 71–79. doi:10.1145/2723872.2723882.

15. Dai M, Wang P, Boyd AD, et al. Evolving gene/transcript definitions significantly alter the interpretation of GeneChip data. Nucleic Acids Res. 2005;33(20):e175. doi:10.1093/nar/gni179.

16. Kopljar I, Gallacher DJ, De Bondt A, et al. Functional and Transcriptional Characterization of Histone Deacetylase Inhibitor-Mediated Cardiac Adverse Effects in Human Induced Pluripotent Stem Cell-Derived Cardiomyocytes. Stem Cells Transl Med. 2016;5(5): 602–612. doi:10.5966/sctm.2015-0279.

17. Karpiński P, Frydecka D, Sąsiadek M. Reduced number of peripheral natural killer cells in schizophrenia but not in bipolar disorder. Brain, Behav. 2016. http://www.sciencedirect.com/science/article/pii/S0889159116300265. Accessed May 31, 2016.

18. Brummelman J, Raeven R, Helm K. Transcriptome signature for dampened Th2 dominance in acellular pertussis vaccine-induced CD4+ T cell responses through TLR4 ligation. Scientific. 2016. http://www.ncbi.nlm.nih.gov/pmc/articles/PMC4846868/. Accessed May 31, 2016.

19. Bilgrau A, Eriksen P, Rasmussen J. GMCM: Unsupervised clustering and meta-analysis using gaussian mixture copula models. J Stat. 2016. https://www.jstatsoft.org/article/view/v070i02/v70i02.pdf. Accessed May 31, 2016.

20. Gandin V, Masvidal L, Cargnello M, Gyenis L. mTORC1 and CK2 coordinate ternary and eIF4F complex assembly. Nature. 2016. http://www.nature.com/ncomms/2016/160404/ncomms11127/full/ncomms11127.html. Accessed May 31, 2016.

21. Killeen A, Diskin M, Morris D. Endometrial gene expression in high-and low-fertility heifers in the late luteal phase of the estrous cycle and a comparison with midluteal gene expression. Physiological. 2016. http://physiolgenomics.physiology.org/content/48/4/306.abstract. Accessed May 31, 2016.

22. Colletti N, Liu H, Gower A, Alekseyev Y. Tlr3 signaling Promotes the induction of Unique human BDca-3 Dendritic cell Populations. Front. 2016. http://www.ncbi.nlm.nih.gov/pmc/articles/PMC4789364/. Accessed May 31, 2016.

23. Lee M, Huang R, Tong W. Discovery of transcriptional targets regulated by nuclear receptors using a probabilistic graphical model. Toxicol Sci. 2015. http://toxsci.oxfordjournals.org/content/early/2015/12/07/toxsci.kfv261.abstract. Accessed May 31, 2016.

24. Troy N, Hollams E, Holt P. Differential gene network analysis for the identification of asthma-associated therapeutic targets in allergen-specific T-helper memory responses. BMC Med. 2016. http://bmcmedgenomics.biomedcentral.com/articles/10.1186/s12920–016–0171-z. Accessed May 31, 2016.

25. Manié E, Popova T, Battistella A. Genomic hallmarks of homologous recombination deficiency in invasive breast carcinomas. J Cancer. 2016. http://onlinelibrary.wiley.com/doi/10.1002/ijc.29829/full. Accessed May 31, 2016.

26. Dekkers B, He H, Hanson J, Willems L. The Arabidopsis DELAY OF GERMINATION 1 gene affects ABSCISIC ACID INSENSITIVE 5 (ABI5) expression and genetically interacts with ABI3 during Arabidopsis. The Plant. 2016. http://onlinelibrary.wiley.com/doi/10.1111/tpj.13118/full. Accessed May 31, 2016.

27. Holt P, Strickland D, Bosco A, Belgrave D. Distinguishing benign from pathologic TH 2 immunity in atopic children. J Allergy. 2015. http://www.sciencedirect.com/science/article/pii/S0091674915013342. Accessed May 31, 2016.

28. Lück S, Westermark P. Circadian mRNA expression: insights from modeling and transcriptomics. Cell Mol Life Sci. 2016. http://link.springer.com/article/10.1007/s00018–015–2072–2. Accessed May 31, 2016.

29. Bosco A, Wiehler S, Proud D. Interferon regulatory factor 7 regulates airway epithelial cell responses to human rhinovirus infection. BMC Genomics. 2016. http://bmcgenomics.biomedcentral.com/articles/10.1186/s12864–016–2405-z. Accessed May 31, 2016.

30. Fauteux F, Hill J, Jaramillo M, Pan Y, Phan S. Computational selection of antibody-drug conjugate targets for breast cancer. Oncotarget. 2015. http://europepmc.org/abstract/med/26700623. Accessed May 31, 2016.

31. Napolitano F, Sirci F, Carrella D, Bernardo D di. Drug-set enrichment analysis: a novel tool to investigate drug mode of action. Bioinformatics. 2016. http://bioinformatics.oxfordjournals.org/content/32/2/235.short. Accessed May 31, 2016.

32. Carroll J, Meyer C, Song J, Li W, Geistlinger T. Genome-wide analysis of estrogen receptor binding sites. Nature. 2006. http://www.nature.com/ng/journal/v38/n11/abs/ng1901.html. Accessed May 31, 2016.

33. Lupien M, Eeckhoute J, Meyer C, Wang Q, Zhang Y. FoxA1 translates epigenetic signatures into enhancer-driven lineage-specific transcription. Cell. 2008. http://www.sciencedirect.com/science/article/pii/S0092867408001189. Accessed May 31, 2016.

34. Wang Q, Li W, Zhang Y, et al. Androgen receptor regulates a distinct transcription program in androgen-independent prostate cancer. Cell. 2009. http://www.sciencedirect.com/science/article/pii/S0092867409005170. Accessed May 31, 2016.

35. Lefterova M, Zhang Y, Steger D. PPARγ and C/EBP factors orchestrate adipocyte biology via adjacent binding on a genome-wide scale. Genes &. 2008. http://genesdev.cshlp.org/content/22/21/2941.short. Accessed May 31, 2016.

36. Tuupanen S, Turunen M, Lehtonen R, Hallikas O. The common colorectal cancer predisposition SNP rs6983267 at chromosome 8q24 confers potential to enhanced Wnt signaling. Nature. 2009. http://www.nature.com/ng/journal/v41/n8/abs/ng.406.html. Accessed May 31, 2016.

37. Obad S, Santos C dos, Petri A, Heidenblad M. Silencing of microRNA families by seed-targeting tiny LNAs. Nature. 2011. http://www.nature.com/ng/journal/v43/n4/abs/ng.786.html. Accessed May 31, 2016.

38. He H, Meyer C, Shin H, Bailey S, Wei G, Wang Q. Nucleosome dynamics define transcriptional enhancers. Nature. 2010. http://www.nature.com/ng/journal/v42/n4/abs/ng.545.html. Accessed May 31, 2016.

39. Ozsolak F, Song J, Liu X, Fisher D. High-throughput mapping of the chromatin structure of human promoters. Nat Biotechnol. 2007. http://www.nature.com/nbt/journal/v25/n2/abs/nbt1279.html. Accessed May 31, 2016.

40. Zuo T, Wang L, Morrison C, Chang X, Zhang H, Li W. FOXP3 is an X-linked breast cancer suppressor gene and an important repressor of the HER-2/ErbB2 oncogene. Cell. 2007. http://www.sciencedirect.com/science/article/pii/S0092867407005454. Accessed May 31, 2016.

41. Enard W, Gehre S, Hammerschmidt K, Hölter S. A humanized version of Foxp2 affects cortico-basal ganglia circuits in mice. Cell. 2009. http://www.sciencedirect.com/science/article/pii/S009286740900378X. Accessed May 31, 2016.

42. Nunez M, Sanchez-Jimenez C, Alcalde J, Izquierdo JM. Long-term reduction of T-cell intracellular antigens reveals a transcriptome associated with extracellular matrix and cell adhesion components. PLoS One. 2014;9(11). doi:10.1371/journal.pone.0113141.

43. Beaulieu-Jones B, Greene C. Continuous Analysis BrainArray: Submission Release Continuous Analysis BrainArray: Submission Release. August 2016. doi:10.5281/zenodo.59892.

44. Duvall P, Matyas S, Glover A. Continuous Integration: Improving Software Quality and Reducing Risk.; 2007. http://portal.acm.org/citation.cfm?id=1406212.

45. Docker. Docker. https://www.docker.com.

46. Beaulieu-Jones BK, Greene CS. Continuous Analysis Example Docker Images. 2016. 10.6084/m9.figshare.3545156.v1.

47. Beaulieu-Jones BK, Greene CS. Continuous Analysis. GitHub repository. https://github.com/greenelab/continuous_analysis. Published 2016.

48. Drone.io. https://drone.io/.

49. Beaulieu-Jones BK. Denoising Autoencoders for Phenotype Stratification (DAPS): Preprint Release. Zenodo. January 2016. doi:10.5281/zenodo.46165.

50. Katoh K, Misawa K, Kuma K, Miyata T. MAFFT: a novel method for rapid multiple sequence alignment based on fast Fourier transform. Nucleic Acids Res. 2002;30(14): 3059–3066. doi:10.1093/nar/gkf436.

51. Rice P, Longden I, Bleasby A, et al. EMBOSS: the European Molecular Biology Open Software Suite. Trends Genet. 2000;16(6): 276–277. doi:10.1016/s0168-9525(00)02024-2.

52. Boj SF, Hwang C-I, Baker LA, et al. Organoid Models of Human and Mouse Ductal Pancreatic Cancer. Cell. 2015;160(1): 324–338. doi:10.1016/j.cell.2014.12.021.

53. Balli D. Using Kallisto for expression analysis of published RNAseq data. https://benchtobioinformatics.wordpress.com/2015/07/10/using-kallistofor-gene-expression-analysis-of-published-rnaseq-data/. Published 2015. Accessed August 1, 2016.

54. Bray NL, Pimentel H, Melsted P, Pachter L. Near-optimal probabilistic RNA-seq quantification. Nat Biotechnol. 2016;34(5): 525–527. doi:10.1038/nbt.3519.

55. Ritchie ME, Phipson B, Wu D, et al. limma powers differential expression analyses for RNA-sequencing and microarray studies. Nucleic Acids Res. 2015;43(7):e47. doi:10.1093/nar/gkv007.

56. Smyth GK. Linear models and empirical bayes methods for assessing differential expression in microarray experiments. Stat Appl Genet Mol Biol. 2004;3:Article3. doi:10.2202/1544-6115.1027.

57. Pimentel HJ, Bray N, Puente S, Melsted P, Pachter L. Differential analysis of RNA-Seq incorporating quantification uncertainty. bioRxiv. 2016. doi:10.1101/058164.

58. Pérez F, Granger BE. {IP}ython: a System for Interactive Scientific Computing. Comput Sci Eng. 2007;9(3): 21–29. doi:10.1109/MCSE.2007.53.

59. Jupyter. http://jupyter.org/. Published 2016. Accessed January 8, 2016.

60. RStudio. RStudio: Integrated development environment for R (Version 0.97.311). J Wildl Manage. 2011;75(8): 1753–1766. doi:10.1002/jwmg.232.

61. Baumer B, Cetinkaya-Rundel M, Bray A, Loi L, Horton NJ. R Markdown: Integrating A Reproducible Analysis Tool into Introductory Statistics. Technol Innov Stat Educ. 2014;8(1): 20. doi:10.5811/westjem.2011.5.6700.

62. Friedrich Leisch. Sweave: Dynamic generation of statistical reports using literate data analysis. Compstat 2002 - Proc Comput Stat. 2002;(69):575–580. doi:10.1.1.20.2737.

63. Beaulieu-Jones BK, Greene CS. Semi-Supervised Learning of the Electronic Health Record with Denoising Autoencoders for Phenotype Stratification. bioRxiv. February 2016. http://biorxiv.org/content/early/2016/02/18/039800.abstract.

64. Souilmi Y, Lancaster AK, Jung J-Y, et al. Scalable and cost-effective NGS genotyping in the cloud. BMC Med Genomics. 2015;8(1): 64. doi:10.1186/s12920-015-0134-9.

